# Bio-medical Big Data Operating System (Bio-OS): An Integrated Data Mining Environment for Data Intensive Scientific Research

**DOI:** 10.1101/2024.10.17.612997

**Authors:** Ji-Long Liu, Li-Guang Yang, Qing-Yu Xiao, Zhao-Qiang Li, Jin-Yu Cheng, Zi-Ru Chen, Jian-Wen Zhou, Xiang Wan, Tsung-Hui Chang, Xin Zhang, Yi-Xue Li

## Abstract

The advent of high throughput sequencing has ushered life science and clinical research into the era of big data, posing significant challenges for reproducibility due to the complexity of data integration and analysis. Although the FAIR principles advocate for the transparent and reliable sharing of scientific data, their implementation remains hampered by technical barriers. The Global Alliance for Genomics and Health (GA4GH) has made strides in standardizing data and tools, yet a comprehensive solution for reproducibility is lacking. In response, we present BioOS, an open source, cloud native Biomedical big data Operating System. This system encapsulates study components data, code, tools, and environments into workspaces, enhancing reproducibility and validation. BioOS employs JSON Schema for machine readability and includes a Hierarchy Hash Mechanism to ensure data integrity. Adhering to GA4GH protocols, BioOS simplifies complex technological implementations, making advanced research tools accessible. Demonstrated through representative workspaces, BioOS fosters seamless research replication, peer review, and editorial evaluation. Its cloud native infrastructure supports dynamic resource allocation, enabling efficient handling of large scale analyses. By integrating AI driven Large Language Models, BioOS enhances user interaction and operational flexibility. As an evolving open source platform, BioOS exemplifies a transformative approach to biomedical research, aligning with FAIR principles and advancing the AI for Science paradigm, thus promoting a more connected, efficient, and impactful research environment.

---

With the advancement of high-throughput biotechnology, life science, and clinical research has entered the era of big data^1,2^. The increased size and complexity of data make analysis reproducible more challenging^3,4,5^. While the FAIR principles promote timely and credible sharing of scientific findings^6,7^, their practical realization is hindered by the intricacies of integrating standard software tools and defining parameters, datasets, and computational environments^8^. The Global Alliance for Genomics and Health (GA4GH) introduced a framework to standardize data, tools, and execution methods^9^. However, comprehensive technical solutions for reproducibility remain elusive. In alignment with GA4GH standards and to further the FAIR principles, we introduce the Bio-OS - a Biomedical big data Operating System - an open-source, cloud-native technology-based, data-intensive research support system in the field of biomedicine (Figure 1). It integrates data resources, operator resources, computational resources, and an application marketplace, providing an AI-driven, four-in-one “brain-like operating system”, which is available at Bio-OS GitHub Repository (https://github.com/Bio-OS).

**Figure 1.**
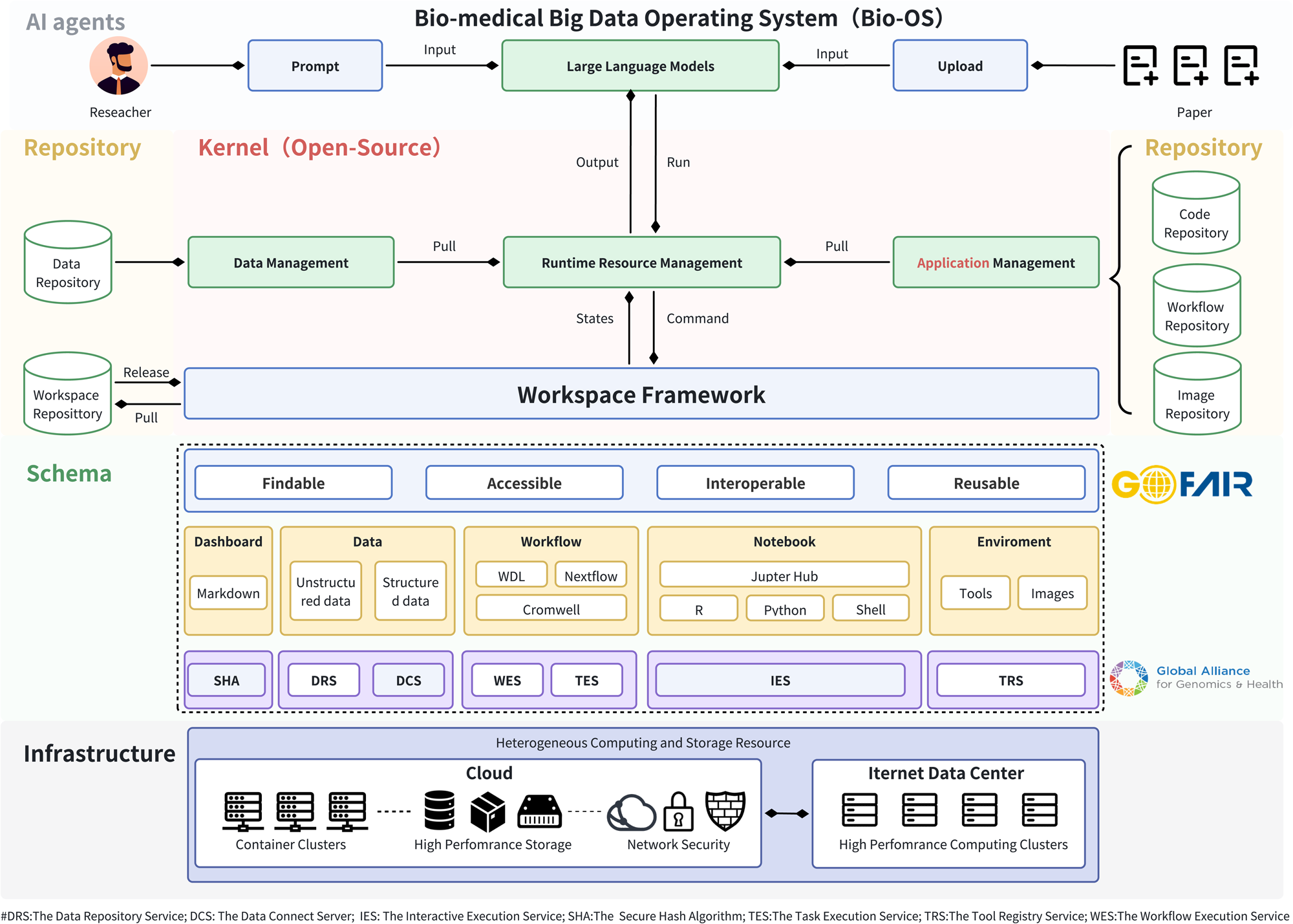
Architecture of Bio-OS. The Bio-OS architecture consists of six modules: Infrastructure, Protocol, Workspace, Kernel, Repository, and AI agents. The infrastructure layer meets the requirements for traditional high-performance computing cluster deployments while also supporting cloud architecture. The Protocol layer ensures compatibility and immutability by supporting GA4GH protocols and incorporating SHA hashing. The Workspace module encapsulates all elements of scientific research to fulfill the FAIR principles. The open-source Kernel layer facilitates decentralized, flexible deployment and resource interoperability. The Repository (coming soon) layer aids in resource sharing and communication. The AI agents (coming soon) layer integrates the latest large language model technologies, enabling interactive analysis through natural language commands.

Historically, researchers predominantly depended on traditional academic publications to broadcast their research, forming the backbone of contemporary scientific paradigms. Yet, the rapid pace of technological progress and evolving research paradigms now challenge the adequacy of literature-driven approach. The ascent of data science and computing has underscored the imperative for sharing raw data, computational tools, and code to bolster study repeatability and transparency. This shift has birthed the “workspace” paradigm. Integral to Bio-OS, the workspace encapsulates every facet—data, code, tools, environment, and descriptions—related to a study as an artefact, empowering peers to reproduce, validate, and build upon initial findings. The Workspace Specification (https://github.com/GBA-BI/bioos-workspace-spec) protocol within Bio-OS offers a structured method for managing research projects, ensuring workspaces are comprehensive and self-descriptive to promote straightforward deposition, dissemination, and reproducibility. Workspace is organized using JSON-Schema and encompasses comprehensive metadata, thus facilitating machine readability and parsing. This brings a lot of benefits. On one hand, machines can interpret various modalities of data, including datasets, workflows, and Docker images, from a Workspace artefact and integrates the multimodal data as input for Al model training to promote Al for Science. On the other hand, Workspace defines clear machine operable application programming interfaces, making it convenient to build Al agents and copilots. A pivotal future enhancement in Bio-OS will be the implementation of the Hierarchy Hash Mechanism (Supplementary Figure 1), which assigns an immutable SHA tag to each workspace and its sub-objects, ensuring integrity and preventing tampering during dissemination. This mechanism utilizes the unique identifiers of GitHub repository URL, Docker image, and MD5 hashes for external files to establish an unalterable record for each component within a workspace artefact. These identifiers are hashed to generate a SHA tag for the workspace, allowing users to verify its integrity and authenticity, a feature that aligns well with blockchain recording and NFT minting concepts.

Bio-OS adheres to GA4GH protocols, such as DRS, WES, TES, and TRS, advancing resource sharing and research collaboration. Bio-OS provides users with the benefits of cutting-edge technological advancements without requiring them to delve into the complexities of the underlying implementations. This ensures that the platform remains accessible, with a low barrier to entry, allowing researchers to leverage new technologies seamlessly. To demonstrate Bio-OS’s proficiency in facilitating research replication and aiding editorial scrutiny, we introduce some representative workspaces (demo workspace, https://mc.miracle.ac.cn/standalone/product/mc/digger/HFV124011/dashboard). Herein, the exhaustive dataset, workflows, and coding framework are amalgamated within a workspace, enhancing the evaluation process for editorial boards, peer reviewers and researchers. The full content of workspaces is directly accessible via a link and distributable as a zip file, enabling import and execution on Bio-OS with a single action.

Diverging from other platforms (Supplementary Table 1) Bio-OS, as an open-source system, exhibits universal accessibility and we are continuously augmenting the repository of open-source material. Concurrently, Bio-OS leverages the essential features of cloud computing, making it more effective in handling large-scale analytical tasks and enabling remote access. With cloud elasticity and scalability, Bio-OS dynamically allocates resources, ensuring continuous research progress even with fluctuating data and computation needs. Furthermore, cloud computing enhances scientific teamwork by facilitating remote data access, application use, and resource sharing, thereby improving research synergy and efficiency. Researchers can not only manage their projects on Bio-OS instance but also directly share their findings with peers by workspace artefacts. This facilitates immediate reproduction of results in different environments with minimal effort. The workspace-centric approach reduces the debugging effort during the process of sharing results. It amplifies the efficiency and reproducibility of research, making the submission and publication of scientific papers more transparent and lucid.

In the realm of life sciences, the integration of big data and artificial intelligence (AI) technologies is pivotal in advancing scientific research. Recognizing the need for advanced tools in scientific research, Bio-OS is evolving to integrate AI-driven Large Language Models (LLMs) features, enhancing user experience and accessibility. The comprehensive knowledge base within Bio-OS mitigates the hallucination issues commonly associated with LLMs, while the tools offer greater operational flexibility for these models (Supplementary Figure 2). In turn, the LLMs provide Bio-OS with improved interactive capabilities. Bio-OS, adhering to the GA4GH framework, combined with LLMs and AI agents, enables AI co-pilots to operate efficiently at an advanced level,perfectly aligning with the AI for Science technological development trend. Collectively, these tools streamline research workflows and data management, promoting cross-platform reproducibility and ensuring comprehensive lifecycle management from data to application. As an open-source system, Bio-OS continues to expand its resources and community for collaborative development, exemplified by the <u>Miracle Cloud instance</u> (https://mc.miracle.ac.cn/login?redirect_uri=%2Fproduct%2Fmc%2Foverview, a comprehensive version tailored for research institutions, with features that will gradually be made available in the open-source version for individual users) that fosters the creation of reproducible, shareable, and reusable research outcomes. More than a collection of technological advancements, Bio-OS represents a transformative approach to biomedical research. By integrating diverse resources into a cohesive, intelligent system, Bio-OS acts as a catalyst for scientific discovery, driving forward the principles of FAIR. Its “brain-like” operation, coupled with cloud-native infrastructure and AI integration, positions Bio-OS as a pioneering force in the digital augmentation of life sciences, leading the way towards a future where research is more connected, efficient, and impactful.

## ACKNOWLEDGMENTS

This work was supported by the CAS Research Fund, Grant No. XDB38050200, the R&D Program of Guangzhou Laboratory, Grant No. SRPG22-001, Self-supporting Program of Guangzhou National Laboratory No. SRPG22-007 and Startup Program of Guangzhou National Laboratory YW-YFYJ0101.

## AUTHOR CONTRIBUTIONS

LG.Y. and YX.L. wrote the manuscript. JL.L. LG.Y. and QY.X and designed the Bio-OS system. ZQ.L., JY.C and JW.Z. contributed to Bio-OS development. QY.X, ZQ.L. and JY.C. wrote the pipeline and automation software. LG.Y. and ZR.C. hosted quantification results. X.W., X.Z. and YX.L. provided scientific leadership and project oversight.

## Supplementary Materials

Supplementary Table 1. Comparison of Bio-OS with other Cloud computing platforms.

Supplementary Figure 1. The basic data structure Workspace in Bio-OS.

**Supplementary Figure 1.**
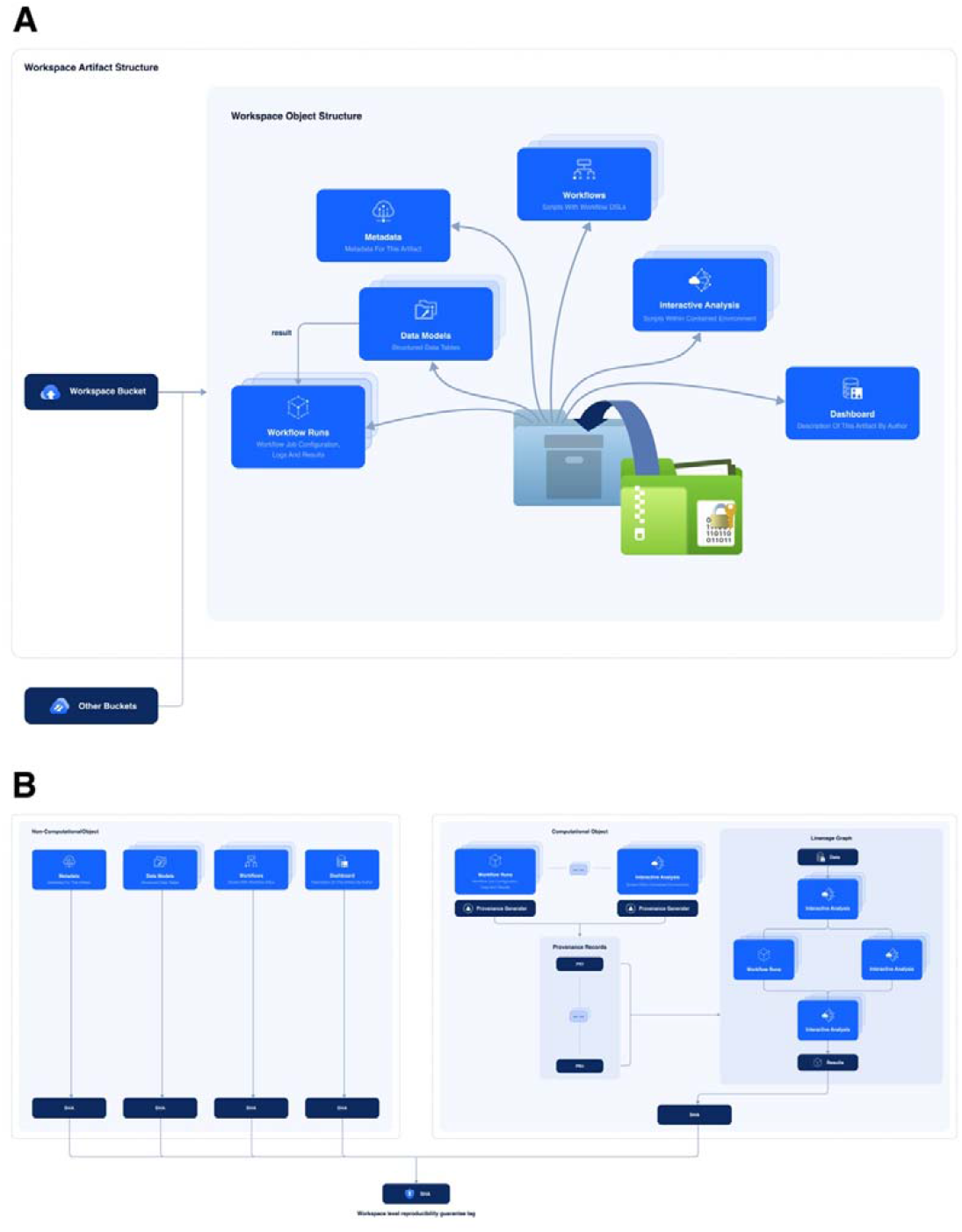
The basic data structure Workspace in Bio-OS. (A) Schematic representation of a Workspace Artifact. It is a lightweight object that encapsulates the entire research project to form one unit for facilitating deposition, dissemination, publication, and reproducibility. (B) Hierarchy Hash Mechanism for Workspace and its sub-objects to achieve immutable Secure Hash Algorithm (SHA) tags. These tags serve as unique identifiers for Workspace during dissemination to ensure integrity and prevent tampering.

Supplementary Figure 2. Bio-OS Agent Demo. By employing large language model methodologies, the tools on Bio-OS facilitate natural language interaction to accomplish analytical tasks.

