## Supplementary figures and images for "Bio-medical Big Data Operating System (Bio-OS): An Integrated Data Mining Environment for Data Intensive Scientific Research"

### Supplementary Figure2

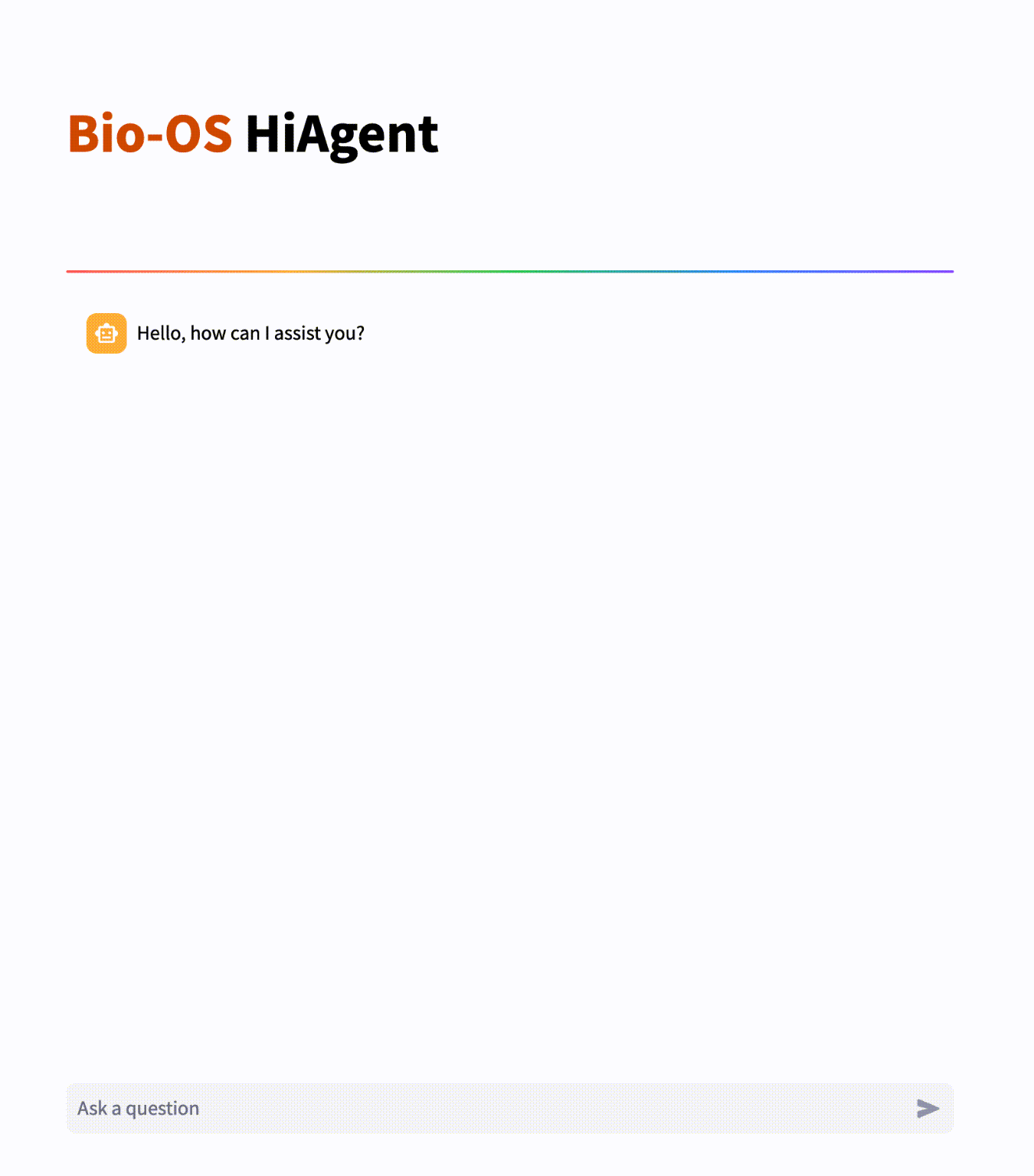
